# A Paradigm Free Regularization Approach to Recover Brain Activation from Functional MRI Data

**DOI:** 10.1101/2021.04.14.438942

**Authors:** Isa Costantini, Rachid Deriche, Samuel Deslauriers-Gauthier

## Abstract

**Context:** Functional MRI is a non-invasive imaging technique that provides an indirect view into brain activity, via the blood-oxygen-level-dependent (BOLD) response. In particular, resting-state fMRI poses challenges to the recovery of brain activity without prior knowledge on the experimental paradigm, as it is the case for task-fMRI. Conventional methods to infer brain activity from the fMRI signals, for example the general linear model (GLM), require the knowledge of the experimental paradigm to define regressors and estimate the contribution of each voxel’s time course to the task. To overcome this limitation, approaches to deconvolve the BOLD response and recover the underlying neural activation without a priori information on the task have been proposed. State-of-the-art techniques, and in particular the Total Activation (TA), formulates the deconvolution as an optimization problem with decoupled spatial and temporal regularization terms. This increases the number of hyperparameters to be set and requires an optimization strategy that alternates between the constraints.

**Approach:** In this work, we propose a paradigm-free regularization algorithm named Paradigm-Free fMRI (PF-fMRI) that is applied on the 4-D fMRI image, acting simultaneously in the 3-D space and 1-D time dimensions. Based on the idea that large image variations should be preserved as they occur during brain activation, whereas small variations considered as noise should be removed, the PF-fMRI applies an anisotropic regularization, thus recovering the location and the duration of brain activation.

**Results:** Using the experimental paradigm as ground truth, the PF-fMRI is validated on synthetic and real task-fMRI data from 51 subjets, and its performance is compared to the TA. Results show higher correlations of the recovered time-courses with the ground truth compared to the TA and lower computational times. In addition, we show that the PF-fMRI recovers activity that agrees with the GLM, without requiring or using any knowledge of the experimental paradigm.

## 1 Introduction

Functional MRI (fMRI) is a non-invasive imaging technique that indirectly probes brain function by providing a measure of the metabolic activity consequent to an increased neural activation. Two experimental setups are commonly used to acquire fMRI data, task-fMRI and resting-state fMRI (rs-fMRI). In the former the subject is asked to follow an experimental paradigm, whereas in the latter the subject is asked to rest in the scanner and not do or think of anything in particular. Standard approaches for the analyses of task-fMRI data are based on the well known general linear model (GLM) adapted by Friston and colleagues in 1998 in the context of fMRI data analyses [1]. This approach requires prior knowledge of the task parameters and timing of events, as well as assumptions about the neural and hemodynamic responses. Therefore, the GLM can be used only for task-fMRI experiments, where the expected stimulus response is given by experimental paradigm. In contrast to task-fMRI, where the focus is on the response to a specific stimulus, rs-fMRI provides insight on brain function in the absence of stimuli. It also allows us to map the brain activity of patients whose condition does not allow them to perform tasks or follow an experimental paradigm. Furthermore, there are also brain activation that cannot be modeled and expected, such as seizures in epileptic patients [2]. For these data, which represent unpredictable brain activity, the GLM approach is not suitable [3].

Data-driven methods have been proposed to analyze images obtained in resting-state, when no information about the occurrence of the activation is available. They include blind source separation approaches such as the independent component analysis (ICA) [4–6], the principal component analysis (PCA) [7, 8], the Temporal Clustering Analysis [9, 10], and clustering methods [11–14]. These methods are of interest if the aim is to group voxels showing the same spatial or temporal features but they cannot be used if the goal is identifying activation at the voxel level. Indeed, they do not consider including any hemodynamic effect. They are also limited by the necessity of choosing a priori the number of components or clusters and by their interpretation [15].

To overcome these limitations, deconvolution approaches have been developed to address the problem of studying and uncovering brain activation hidden within fMRI time series at the voxel level. Functional MRI deconvolution was introduced by Glover in 1999 [16], who investigated the performance of Wiener deconvolution for deblurring the fMRI response and reduce image distortions. This approach resulted in smooth recovered activation [2] and required an independent measurement of the noise spectral density [17]. Gitelman et al. [17] developed an approach based on linear deconvolution and modeled the interplay between areas as effects of psycho-physiological interactions. In addition, dynamical filter methods, such as Kalman and Bayesian filtering, and Local Linearization filters have been developed and applied to fMRI [18–20]. However, because these approaches are based on non-linear models in continuous time, they are limited by the high computational cost and convenient only for the analysis of localized regions of interest (ROIs) [2]. Other approaches make spatial and/or temporal assumptions on the underlying signals, thus adding priors in the optimization problems. In particular, sparse regularization on the recovered activation maps were exploited to force to zero the weights of regressors that did not contribute to the activation [21–23]. L1-norm regularization approaches have also been developed to exploit sparse temporal features of the hidden neural activation. This was done by means of the majorization-minimization of a cost function to find an optimal solution to the inverse problem [24]. Caballero Gaudes et al. developed a ridge-regression regularization by minimizing both the variance of the residuals and the power of the resulting estimate of the input signal representing the brain activity [15] and a sparse regression [25] by assuming short neuronal activation. Temporal regularized optimization problems based on wavelets were also explored [26]. These methods exploit the temporal features of the hemodynamic response function (HRF) [2] attempting to deconvolve it from the fMRI data, rather than using any information on the timing of the neural activation which translate on the blood-oxygen-level-dependent (BOLD) response. Recently, by supposing the brain activates in constant blocks, Karahanoğlu and colleagues [2], later revisited by Farouj et al. [27], developed a deconvolution approach that involves both spatial and temporal regularization called Total Activation (TA). These approaches split the optimization problem into two decoupled spatial and temporal regularization which increases the number of parameters to be set and requires the solver to alternate between the constraints. The temporal regularization of the latter was then improved by Costantini et al. [28] by proposing a joint approach using the least angle regression (LARS) algorithm and the L-curve that overcame the limitation of having to choose a priori the regularization parameter, and reduced the computation time.

Starting from the idea that large image variations should be preserved as they occur during a brain activation, whereas small variations considered as noise should be smoothed, we propose a novel approach, based on partial differential equations (PDEs), named Paradigm-Free fMRI (PF-fMRI). The PF-fMRI applies an anisotropic diffusion process whose diffusivity is steered by derivatives of the evolving image, to smooth the fMRI image and simultaneously enhance important features such as spatial edges and temporal functional activation. Regularization methods have been enriched by the use of non-linear PDEs in several contexts for the last 30 years. Firstly applied to physics and fluid mechanics, it has been shown that non-linear PDEs allow smoothing the data while preserving large global features, such as discontinuities of the signal [29], which can be found, for example, in image contours and corners [30]. This approach is rooted on the isotropic diffusion equation, i.e. heat flow, and has subsequently been extended to other theoretical contributions. Among them there are the anisotropic smoothing [31, 32] and the PDEs-based gradient descent used to solve energy functionals minimizations [33–37]. The pioneering work that employed anisotropic diffusion PDEs for the restoration of noisy and blurred digital data was proposed by Perona and Malik in 1990 [38] overcoming the limitations associated to linear filtering approaches [39]. To date, PDEs-based regularization algorithms have been applied to 2-D scalar images [31, 36, 38, 40] and vector-valued images [39].

The PF-fMRI proposed in this work has been conceived for the geometrical regularization of 4-D fMRI images (3-D space × 1-D time) based on PDEs. The PF-fMRI acts concurrently in space and time thus overcoming the limitation of previous deconvolution approaches which consider the two problems of spatial and temporal regularization as decoupled processes. In our method, the regularization flow is performed according to the time and to the local geometry of the image to evaluate the presence of an edge and its local strength.

The rest of the paper is organized as follows. We first introduce the PDEs regularization theoretical framework, we illustrate the mathematical problem and how we solved it. Next, we validate the PF-fMRI on task-fMRI data from 51 subjects using the experimental paradigm as ground truth and positively compare its performance to the TA approach. Finally, the advantage of our approach of not requiring any knowledge of the experimental paradigm is shown by recovering activity that agrees with the GLM that uses and need this a-priori knowledge.

## 2 Theory

Inspired by the physics of fluids, many authors assimilated the process of image regularization with the diffusion of chemical concentrations and proposed to apply the following diffusion PDE process [29–31, 39, 41]

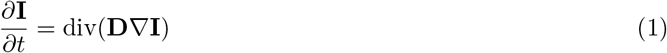

where **I** is the input image, ∇ is the gradient operator, *t* is the time, div(·) is the divergence operator and

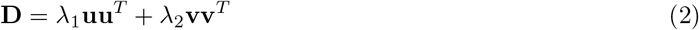

is the diffusion tensor of the image **I**, also called structure tensor [29, 42, 43]. The diffusion tensor **D** is a symmetric and positive definite matrix, and has *λ*_1_, *λ*_2_ as positive eigenvalues and **u, v** as corresponding orthogonal eigenvectors that drive the regularization process; the amount of diffusion in the directions **u** and **v** will be weighted by *λ*_1_ and *λ*_2_, respectively. PDEs smooth the image at each step with a notion of scale-space [38, 40, 44]; at each iteration, the image is smoothed and fine-scale properties, such as noise in our case, are gradually suppressed. To apply the diffusion process to fMRI data, Eq. (1) will need to be reformulated to handle 4-D images. Let us define a scalar-valued image as a function **I** : Ω⊂ℝ^4^, where Ωis the domain of the 4-D (3-D space × 1-D time) image and let us assume Neumann boundary conditions on *δ*Ω, specifying the values in which the derivative of the solution is applied within the boundary of the domain. Let us now define a structure tensor **D** as a 4 × 4 symmetric and positive-definite matrix. By definition, **D** has four positive eigenvalues (*λ*_1_ ≥ *λ*_2_ ≥ *λ*_3_ ≥ *λ*_4_ ≥ 0) and their associated four orthogonal eigenvectors (***θ***_1_, ***θ***_2_, ***θ***_3_ and ***θ***_4_) explain the distribution and orientation of the gradient ∇**I** = (*I*_*x*_, *I*_*y*_, *I*_*z*_, *I*_*t*_) of the image **I** in a given neighborhood. A structure tensor can distinguish between anisotropic and isotropic diffusion. If *λ*_1_ *>> λ*_2_, *λ*_3_ and *λ*_4_, the structure tensor has a principal orientation (in this case ***θ***_1_) and the diffusion is anisotropic. It can be represented with an ellipsoid oriented along ***θ***_1_. On the other hand, if *λ*_1_ ≈*λ*_2_ ≈*λ*_3_ ≈*λ*_4_, the structure tensor is not oriented in a main direction and ***θ***_1_, ***θ***_2_, ***θ***_3_ and ***θ***_4_ are eigenvectors of **D** with equal weight. In this case, the diffusion is isotropic and the structure tensor can be represented with a sphere.

Inspired by the physical process of diffusion, we link the diffusion to fMRI image regularization and we propose the PF-fMRI for the enhancement of coherent structures found in fMRI data. The PF-fMRI recovers brain activation and smooths small variations while preserving large variations via a regularization that is applied on the 4-D fMRI image, acting simultaneously in the 3-D space and the 1-D time dimensions. Using this approach corresponds to performing minimization of image variations as well as a blind image deconvolution. To do this, we propose a regularization process based on a gradient descent computed with PDEs, such that

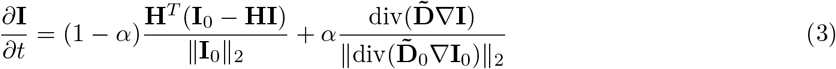

where the term on the left, *∂***I***/∂t*, is the regularization flow, the first term on the right is the data fitting term, also called fidelity term, that prevents the solution from straying far from the input data, and the second term on the right minimizes image variations. The parameter *α* ∈ [0, 1] is the user-defined regularization parameter that balances the fidelity and the regularization terms. Starting from the initial image **I**_0_, the restored image **I** is regularized as *t* increases reducing noise and extracting coherent space and time variations. At the same time, no new structures are introduced in the image [36].

Going more into the details of the fidelity term

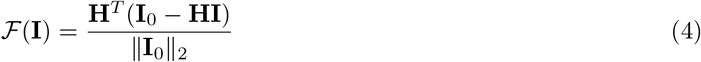

where **I**_0_ and **I** are the original and the regularized image respectively, ∥**I**_0_ ∥_2_ in the denominator is the normalization factor, **H** is the HRF [26] operator, and **H**^*T*^ is its transpose. The multiplication of **H**^*T*^ with (**I**_0_ − **HI**) corresponds to a correlation and can be implemented via convolution with the time-reversed HRF. Note that this product is computed only along the time dimension. The HRF considered in this work is the linearized time-HRF operator proposed by Khalidov et al. [26, 45].

The regularization term is defined as

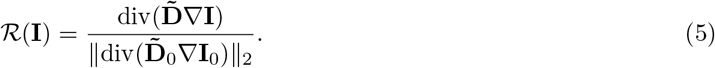

where 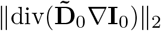 is the normalization term and 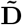 is the regularization tensor, distinct from the structure tensor **D**. In order to elucidate the regularization term, let us start by the definition of the operator

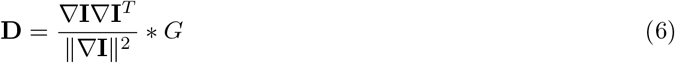

that is the 4-D structure tensor of **I** smoothed by the gaussian kernel *G* with standard deviation *σ*_*G*_, via the convolution operator *. The matrix **D** being the diffusion tensor of the image **I**, its eigendecomposition gives a set of eigenvalues and eigenvectors such that, if the gradient in one direction is large, the eigenvalue associated to that direction is large, whereas the eigenvalues associated to the other three directions are relatively small. Since we are processing fMRI images with the aim of identifying activation and contours that occur concomitant to a large gradient in a certain direction, we aim at reversing the diffusion process, therefore at reversing the effect of **D** into 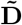 to enhance and simultaneously simplify coherent structures of the fMRI image. Here, we propose to compute the operator 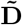 by modifying the eigenvalues of the operator **D** in Eq. (6). Specifically, we defined the directions of the image variations by an eigendecomposition of **D** such that

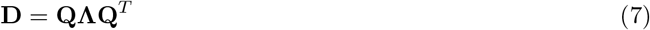

where **Q** contains the orthogonal eigenvectors (***θ***_1_, ***θ***_2_, ***θ***_3_, ***θ***_4_) of **D** and **Λ** contains their associated eigenvalues (*λ*_1_ ≥ *λ*_2_ ≥ *λ*_3_ ≥ *λ*_4_). We then recomputed the matrix

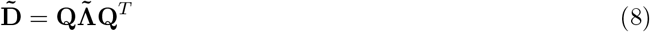

where 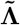 is a diagonal matrix with entries 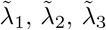, and 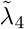 such that for each voxel the highest eigenvalue is given by

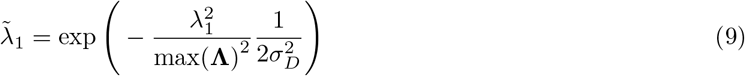

and the other eigenvalues are 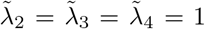. The rationale is that, if *λ*_1_ is large, the current voxel may be located on a edge or activation and the diffusion tensor 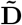 is steered to be anisotropic, by setting 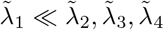. Since we aim at performing a smoothing only along the other three directions to smooth preferably along the coherence directions, the three eigenvalues *λ*_2_ ≈*λ*_3_ ≈*λ*_4_ are set to 1. On the other hand, if *λ*_1_ is small, the diffusion will be isotropic in the four directions because 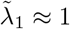 and 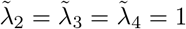 Using the function in Eq. (9) corresponds to reassigning to each voxel different eigenvalues constituting the matrix 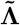, before recomputing the operator 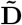 as in Eq. (8). In fact, if *λ*_1_*/*max(**Λ**) is large, the highest eigenvalue 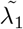 of the considered voxel will tend to zero. This steers the geometrical regularization to be anisotropic, because the smoothing will apply equally in the remaining three directions but it will be negligible in the normal to the detected contour. Otherwise, if *λ*_1_*/*max(**Λ**) is small, the greatest eigenvalue 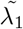 will tend to 1 and this leads to an isotropic regularization almost in all the four directions (x, y, z, t). In both cases indeed 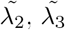, 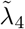are set to 1. This procedure is applied to each voxel of the entire 4-D image such that at each iteration the image **I** computed in Eq. (3) is removed from the image at the previous iteration. In this way, supposing the brain activates in constant blocks, we regularized the image together in space and time. We were able to keep large image variations occurring during brain activation or spatial edges, and to gradually remove small variations, corresponding to noise, while conserving and enhancing coherent structures of the fMRI image. In the following sessions we will explain how we validated the PF-fMRI on phantom and real data.

## 3 Methods

### Simulation of fMRI Data

To reproduce the acquired fMRI signals, for each voxel *v*, we modeled the activity-inducing signal as a boxcar function

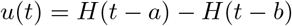

where *H*(*t*) is the Heaviside step function. We added noise to *u*(*t*) representing the random intrinsic electrical fluctuations within neuronal networks that are not associated with encoding a response to internal or external stimuli. To do this, we corrupted the activity-inducing signal *u*(*t*) with an additive random gaussian noise with zero mean and standard deviation *σ*_*m*_ that we called “model noise” *ϵ*_*m*_. The noisy activity-inducing signal is

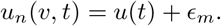

We modeled the activity-related signal *x*(*t*), consequent to the neural activation as the convolution of *u*_*n*_(*t*) with the HRF, modeled as the linear time-invariant system *h*(*t*) [26]:

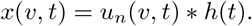

Real time series acquired using the fMRI technique are corrupted by different kinds of noise and artifacts given by mechanisms that do not reflect any neurophysiological function, such as heart rate, respiratory fluctuations, motion artifacts, thermal noise and scanner drifts [46]. For this reason we added noise to *x*(*t*) thus obtaining the acquired fMRI signals

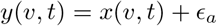

where *ϵ*_*a*_ is the additive random gaussian noise with zero mean and standard deviation *σ*_*a*_. A scheme representing the model of the phantom fMRI data is shown in Figure 1.

**Figure 1.**
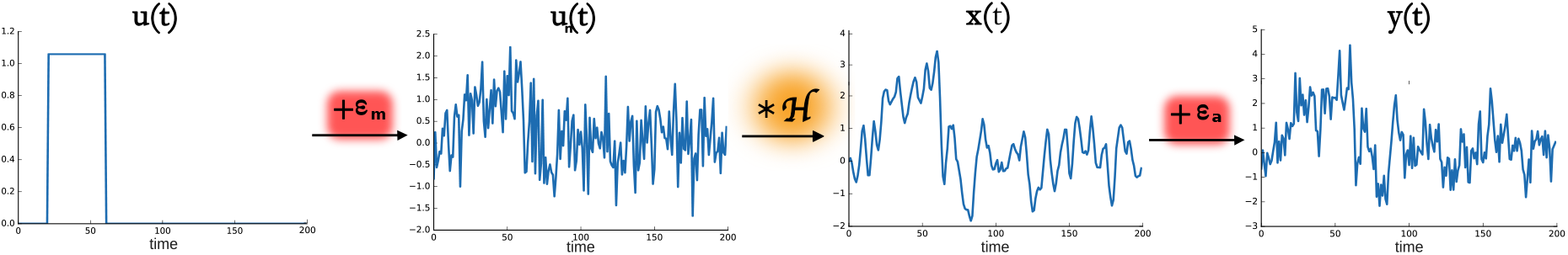
fMRI model. From left to right, *u*(*t*) is the activity inducing signal, that represents the neural activation as a piece-wise constant signal. To this signal a model noise *ϵ*_*m*_ with a gaussian distribution was added thus leading to the signal *u*_*n*_(*t*), that was then convolved with the hemodynamic response function operator ℋ, thus obtaining *x*(*t*), also called activity-related signal. Adding the noise *ϵ*_*a*_ to the signal *x*(*t*) gives the simulated acquired fMRI data denoted as *y*(*t*).

### Validation on synthetic data

To test and validate the PF-fMRI, we scaled a 3-D activation map computed with the FMRIB Software Library (FSL ^1^) Physics-Oriented Simulated Scanner for Understanding MRI (POSSUM ^2^) in the range [0, 3], with a 2-mm isotropic resolution (Figure 2.a). We multiplied it by a piece-wise constant signal *u*(*t*) of 100 s, with one onset of 40 s, from 20 s to 60 s (Figure 2.b). Starting from *u*(*t*) we simulated the acquired fMRI time-courses *y*(*t*) as explained in the previous section. We tested the PF-fMRI on several simulated images obtained by adding different amount of noise for each experiment. We regularized the whole image using the PF-fMRI as shown in Section 2, and we recovered the voxel-wise activity-inducing signals 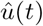. To evaluate the results, we computed the root of the mean square errors (MSEs) and standard deviation (STD) and the Pearson correlation (r) and its STD between *u*(*t*) and 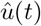 averaged among the voxels belonging to the gray matter (GM). We compared our results with those obtained using the TA approach [27], implemented in the TA toolbox^3^ which constrains the computation only on the GM voxels.

**Figure 2.**
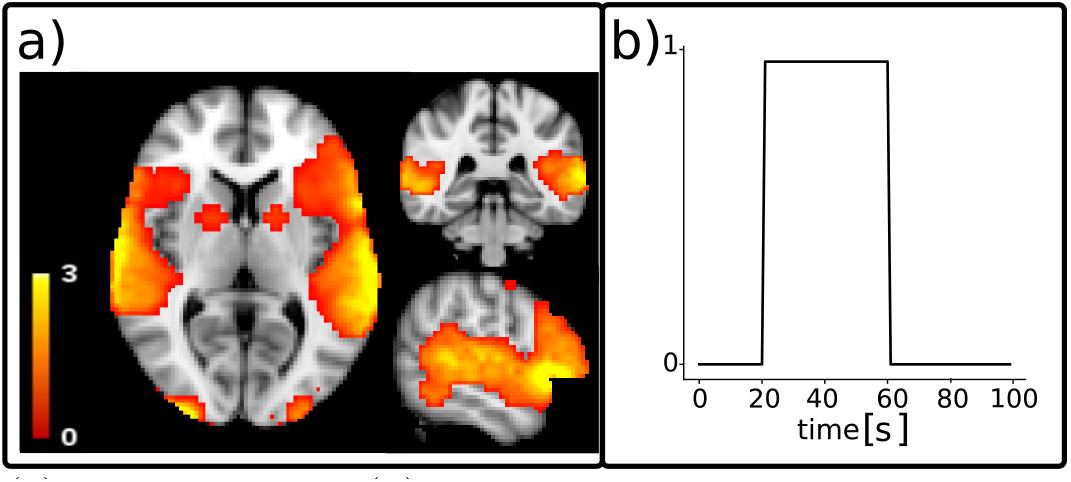
Spatial map (a) and time series (b) of the activation considered as ground truth for functional MRI simulated data. The time course of the activation *u*(*t*) was simulated with a repetition time of 1 s.

### Validation on real experimental data

In addition to validating the PF-fMRI on simulated data, we evaluated its performance on real data. Although our approach was conceived to be applied to rs-fMRI data, or in any situation where no experimental paradigm is given, we test it here on task-fMRI data. This testing strategy provides as with a ground truth i.e. the task timing, which can be used to assess the performance of the algorithm.

The study was conducted on the motor task-fMRI data from 51 subjects from the Human Connectome Project (HCP) database [47]. The data underwent a minimal pre-processing pipeline [48] which includes: correction of gradient-non linearity-induced distortions; registration of each image frame to the signal-band reference image to achieve motion correction; phase-encoding distortion correction; EPI image distortion correction; registration of the fMRI volumes to the structural data; coregistration of the fMRI data to the Montreal Neurological Institute (MNI) space; masking and fMRI image intensity normalization to the 4-D whole global mean of 10^3^. As additional pre-processing steps, each voxels’ time course was detrended to remove linear trends and normalized to 0 mean and unit standard deviation. The motor task is initiated by a visual cue followed by the movement of the left and right foot, the left and right hand, and the tongue. The tasks starting points were considered equal for each subject and inter-subjects differences of the order of milliseconds were neglected.

After applying the PF-fMRI on the entire brain images of each subject, we recovered the reconstructed activity-inducing signals 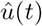 without prior knowledge on the onset/offset times and location of the evoked stimuli. The regularization parameter *α* was set experimentally to 0.9997, *σ*_*G*_ was set to 1, *σ*_*D*_ was set to 0.2 and we computed up to 40 iterations.

To highlight the ability of the PF-fMRI to recover brain activation without knowledge of the experimental paradigm, we qualitatively compared brain regions recovered using the PF-fMRI to those recovered using the GLM as implemented in the FSL library. The GLM model requires as input the exact occurrence of the tasks that the subject is asked to perform. We run the PF-fMRI on the whole brain thus blindly recovering brain activation, without prior knowledge on the intervals of the evoked stimuli. To estimate the results obtained using the PF-fMRI, we computed the voxel-wise correlation maps, by estimating the Pearson correlation coefficient (*r*) between the recovered activation and the 5 tasks. The tasks were simulated as a piece-wise constant signal with unit amplitude when the subject is performing the task and zeros elsewhere. As for the GLM, we included the 5 tasks in a design matrix and we estimated the regressors weights with FSL. Results showing differences and similarities of both approaches were qualitatively assessed.

Subsequently, we quantitatively compared the results obtained using the PF-fMRI with the ones given by the TA, on the sample data composed by 51 subjects. We first defined 4 ROIs located in brain regions that are involved in the 5 considered tasks. The ROI related to the tongue was bilateral whereas for the hands and the feet we defined separate ROIs for the left and the right side of the brain. To do this, we started by defining the ROIs from the work proposed by Roux et al. in 2018 [49], who mapped the somatosensory homunculus MNI coordinates using the electrostimulation. For each coordinate center, we built a spherical 3mm-radius ROI and we grouped the multiple ROIs related to each task into a unique ROI. Coordinates’ centers are shown in Table 1.

**Table 1.**
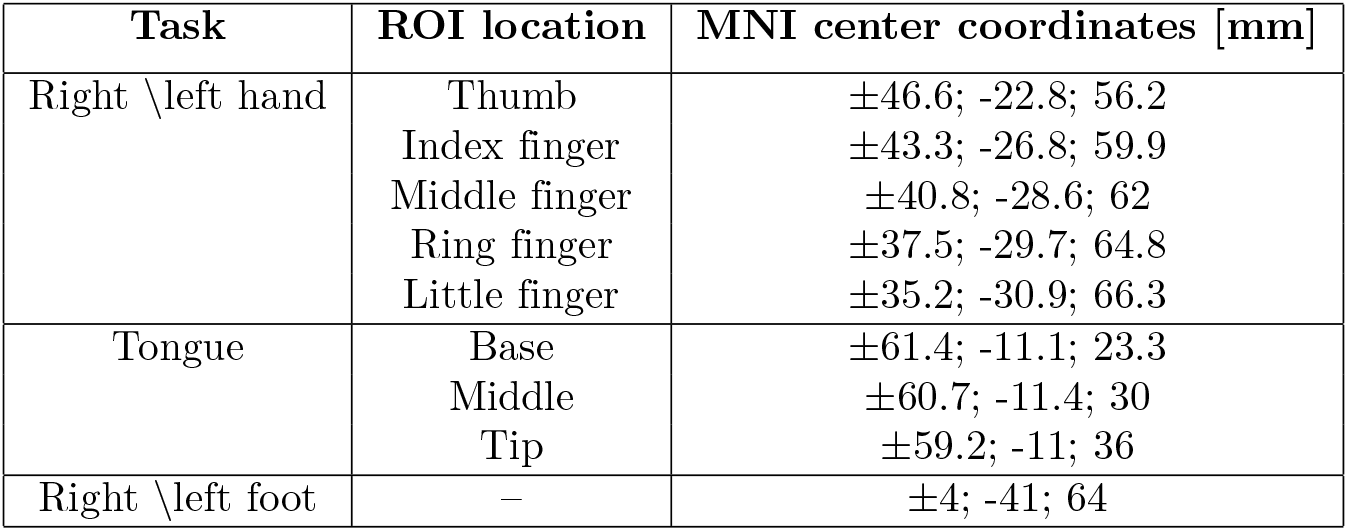
MNI coordinates centers of the brain areas found to respond to the somatosensory stimulation. The coordinates, adapted by Roux et al. [49] are expressed in MNI standard space.

After defining the ROIs, similarly to the comparison between the PF-fMRI and the GLM, we computed the whole-brain voxel-wise correlation maps between the time course related to each task and the recovered activity-inducing signals 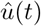 obtained with the PF-fMRI and the TA. For each subject, we first computed the average of the Pearson correlation coefficients (*r*) inside the GM-masked ROIs and then calculated the mean and the standard deviation of these averaged correlation values across the 51 subjects belonging to the sample data. Furthermore, to show that the PF-fMRI is able to differentiate between a region that is active and one that is not, the time courses 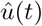 of one representative subject (100307), were averaged in two ROIs of 6 × 6 × 6 *mm*^3^: one that is expected to be active during the task, and one located in a brain area which is not involved in the task. We selected the task related to the tongue, and we chose one ROI centered in the Brodmann Area 4p (rBA4p; MNI coordinates: 62, -14, 30) that is activated during a tongue motor task, and another centered in the primary auditory cortex (TE1.2; MNI coordinates: 56, 4, 10) [50], that is not involved in the tongue movement. The Pearson correlations (*r*) were computed between the tongue activation and the recovered 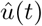 for each voxel, and then averaged among the voxels belonging to the two GM-masked ROIs. Again the tongue activation was simulated as a piece-wise constant signal with unit amplitude during the task and zeros elsewhere. We compared results obtained using the PF-fMRI with the ones obtained using the TA toolbox.

Finally, we applied the PF-fMRI on the rs-fMRI image of one subject (100307) from the HCP database. The data were acquired with a SIEMENS MAGNETOM Connectome Syngo MR D11 using a gradient-echo EPI sequence (TR = 720 ms; TE = 33.1 ms; flip angle = 52°; FOV = 208×180 mm; slice thickness 2.0 mm; number of slices = 72; 2.0 mm isotropic voxels; multiband factor = 8). The subject was asked to lay in the scanner without thinking about anything in particular. The number of acquired frames were 1200 and the duration of the acquisition was 14:33 min. In the case of rs-fMRI data, the task paradigm is unavailable since the subject does not perform any task in the scanner. The data underwent the same minimal preprocessing of the task fMRI data as proposed in the HCP pipeline [48]. In addition, the time series were detrended to remove linear drifts and normalized to zero mean and unit standard deviation. We applied the PF-fMRI algorithm on the entire rs-fMRI sample and we observed the dynamics of the recovered activation maps across time.

## 4 Results

### Validation on synthetic data

Figure 3 shows examples of regularized spatial maps (Figure 3.a) and time series (Figure 3.b) using the PF-fMRI 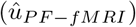 and the TA 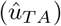. Both approaches do not require any prior knowledge of the paradigm timing. The regularized spatial maps in Figure 3.a represented in the axial plane show how the regularized fMRI image recovered with the PF-fMRI 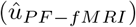 is closer to the ground truth (*u*) in terms of signal amplitude with respect to those obtained using the TA 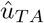. This is verified for different peak-SNRs (pSNRs), i.e. 6.54 dB, 5.99 dB, 5.9 dB and 3.93 dB. In Figure 3.b we show examples of regularized time courses and that we recover an amplitude closer to the ground truth when compared to the method implemented in the TA tool. We also show smoother recovered signals when compared to the TA.

**Figure 3.**
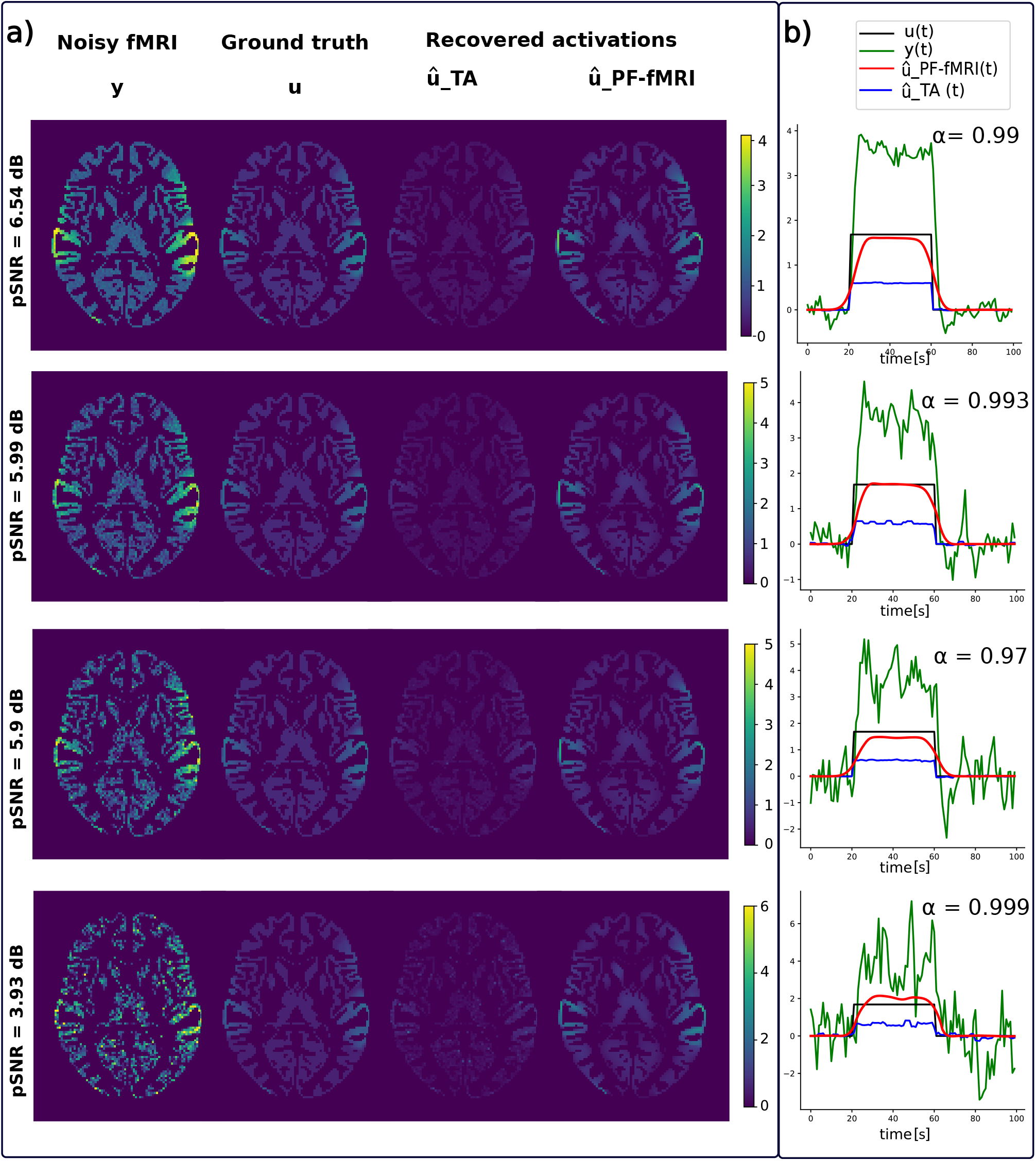
**(a)** From left to right: spatial maps of the simulated fMRI image *y*, ground truth activation *u*, recovered activation using the TA approach 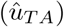 and our approach 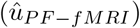. Each row corresponds to a different peak-SNR (pSNR): 6.54 dB, 5.99 dB, 5.9 dB, 3.93 dB from the top to the bottom. **(b)** Reconstructed time series 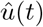 obtained with our approach (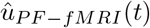, red) and the TA approach (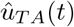, blue) superimposed on the activation (*u*(*t*), black) and fMRI signal (*y*(*t*), green).

Figure 4 shows that the roots of MSEs± STDs computed between the simulated activation *u*(*t*) and the recovered one 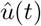 change for different pSNRs. We show lower errors with lower standard deviations than the ones obtained using TA.

**Figure 4.**
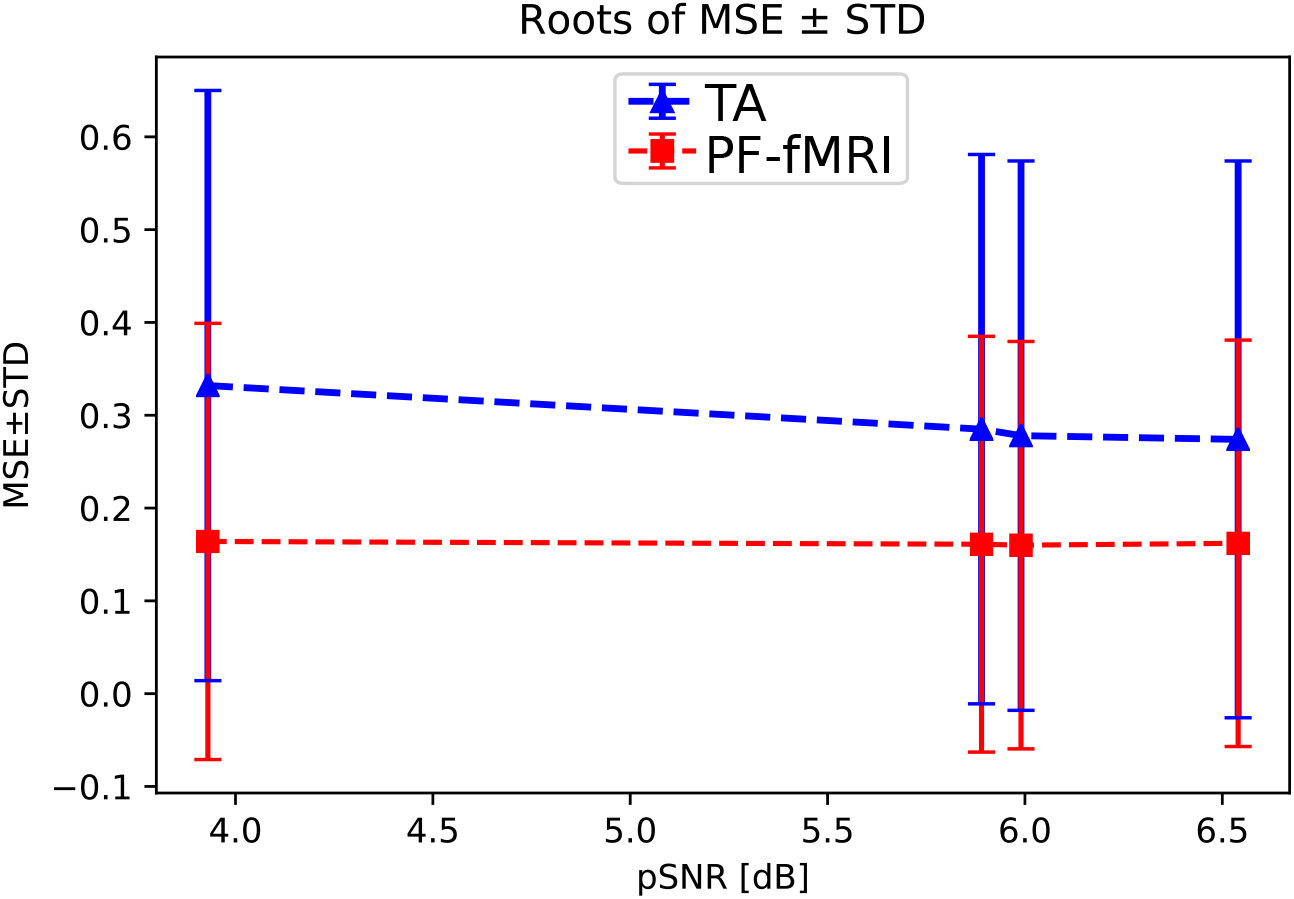
The graph shows, for different pSNRs, the roots of the mean squared errors (MSEs) and standard deviations (STDs) between *u*(*t*) and 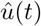 averaged among the voxels belonging to the gray matter (GM).

Figure 5 shows that the activation recovered with the PF-fMRI is more correlated with the ground truth (*r*≈1), for different pSNRs. Whereas the results obtained with TA are more sensitive to noise and show better performances for less noisy data: the mean correlation increases according to the pSNR while the standard deviation decreases.

**Figure 5.**
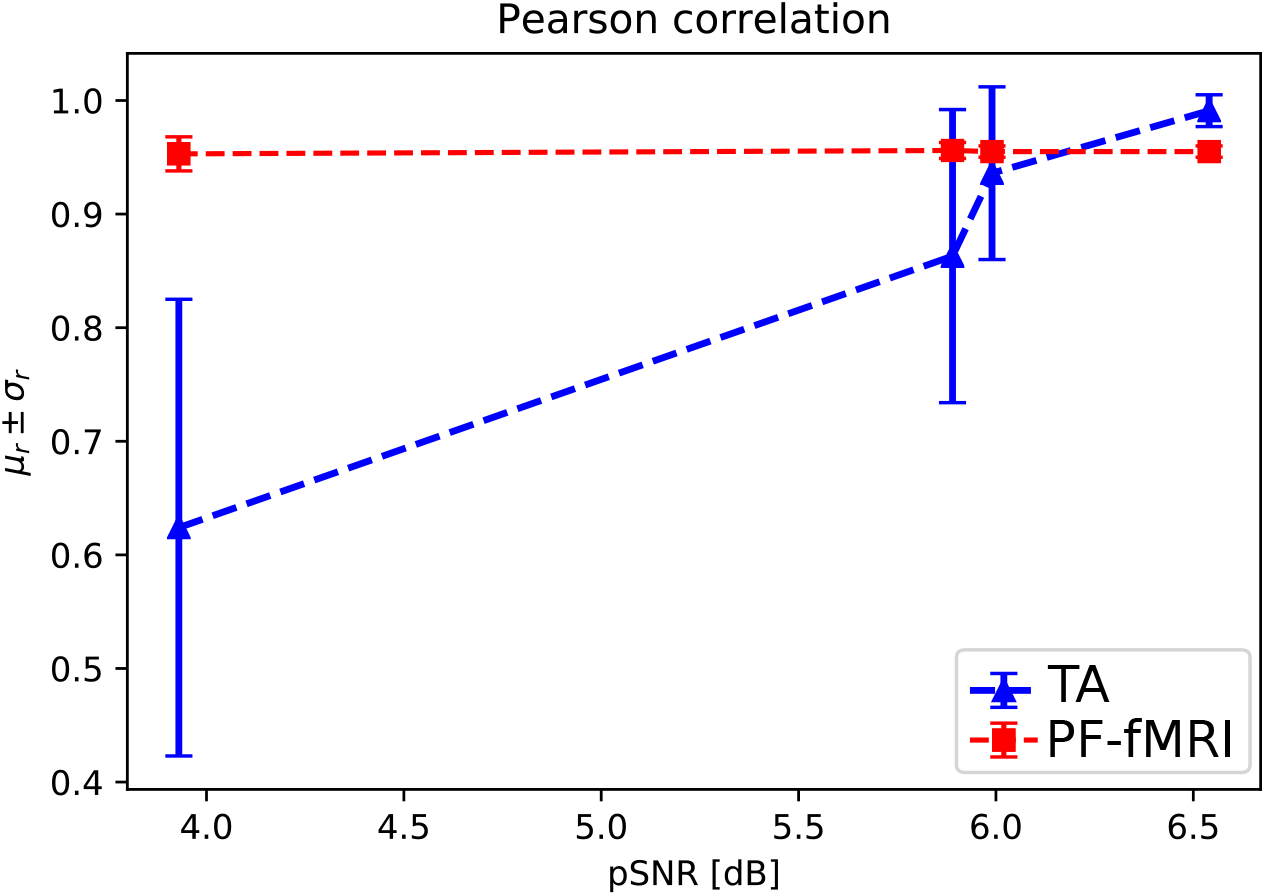
The graph shows, for different peak-SNRs (pSNRs), the mean Pearson correlation coefficients (*µ*_*r*_) and related standard deviation (*σ*_*r*_) computed between *u*(*t*) and 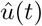 and averaged among the voxels belonging to the GM.

### Validation on real data

Figure 6 shows the regularization tensors 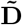 (Eq. 8) in a coronal brain slice for a representative subject of the HCP database. Note that the images refers to the spatial maps, and the temporal dimension is not represented. The presence of ellipsoids oriented in different directions rather than spheres shows the anisotropic nature of the regularization.

**Figure 6.**
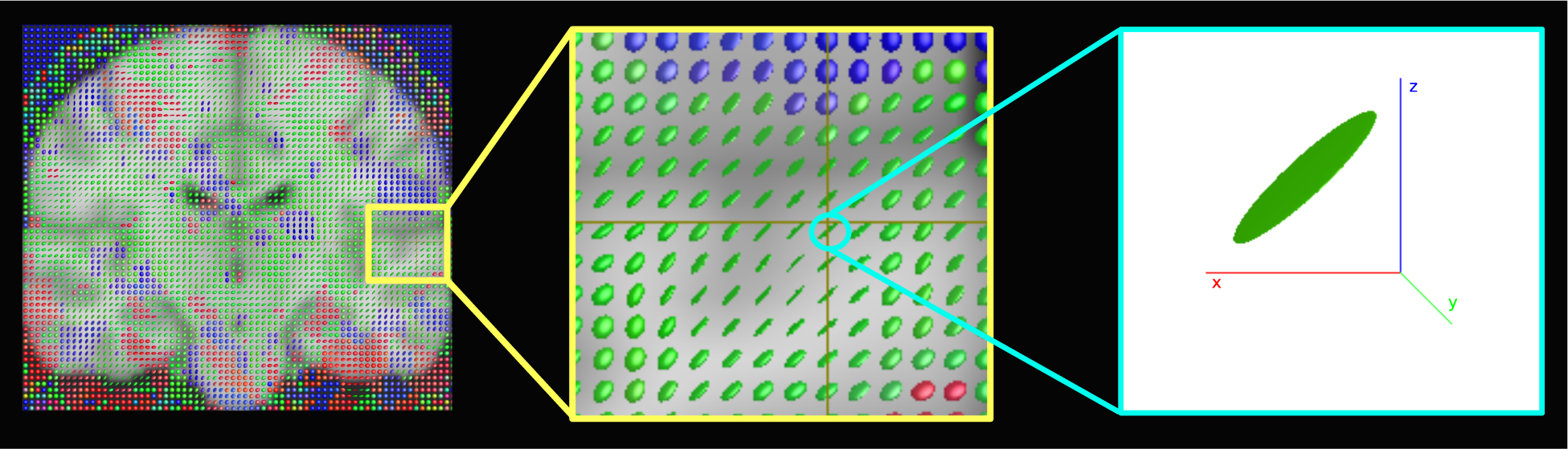
Representation of the structure tensors computed with the PF-fMRI. The images show for a 3-D spatial map, superimposed to the standard MNI template, how the structure tensors look like ellipsoids or spheres meaning that the geometric regularization is applied anisotropically. Images were made using MRview [51].

As for the real data analyses, and specifically the comparison between the PF-fMRI and the GLM, we show that the correlation maps related to each task computed with the PF-fMRI were well overlapped to the values of the regressors coefficients obtained using the GLM as shown for one illustrative subject (100307) in Figure 7. The GLM shows results that follow the GM, while the activation found with the PF-fMRI, that again were performed across the whole brain, and not masked with the GM mask, covers also voxels across the white matter. Interestingly, the found activation overlaps the areas found to be active in the motor homunculus brain [52].

**Figure 7.**
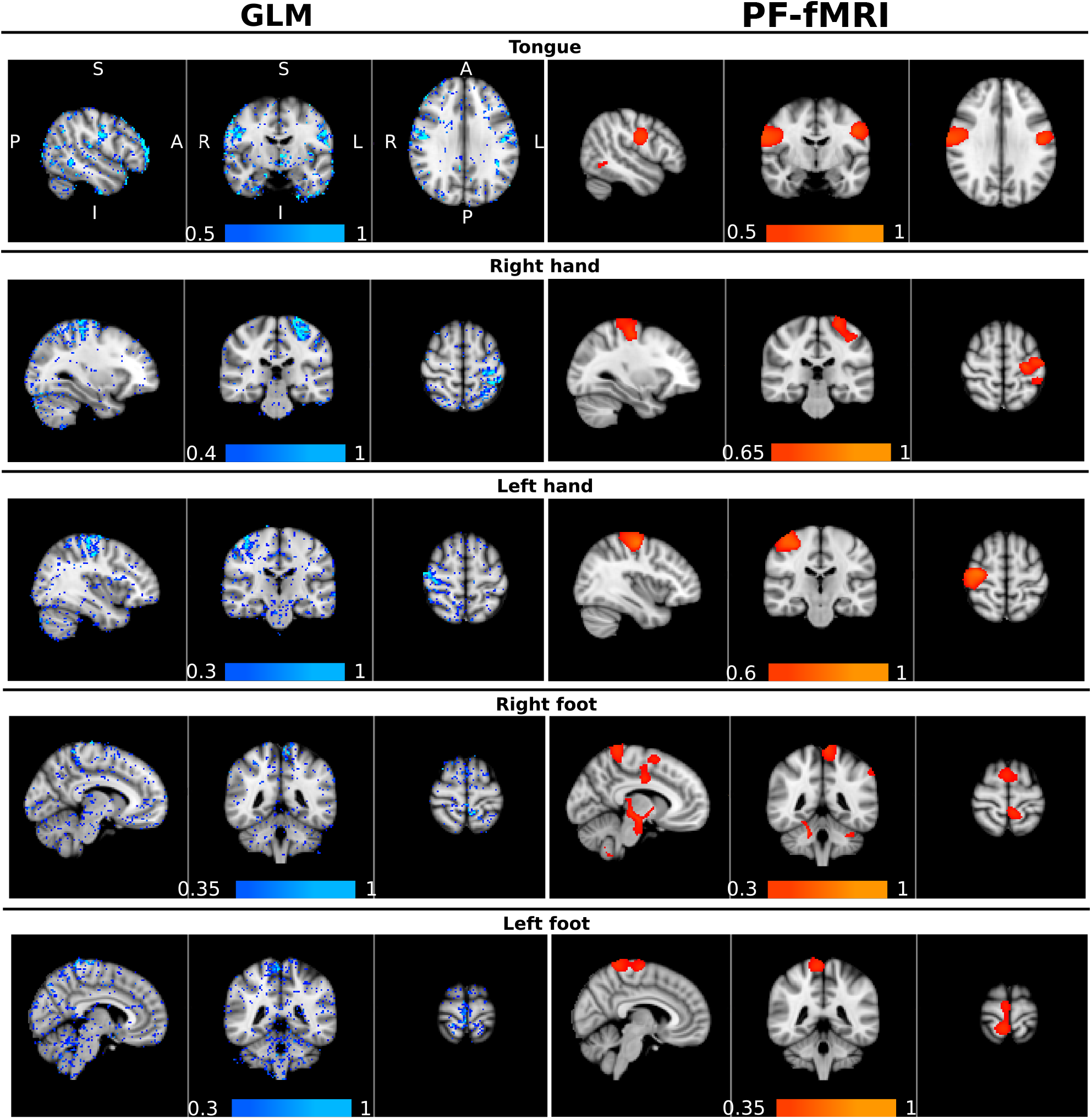
Qualitative comparison between the GLM and the PF-fMRI. On the left column, in a blue-lightblue color-map, superimposed to the standard MNI template, the *β*-regressors map obtained using the GLM implemented in FSL. On the right column, in a red-yellow color-map, the whole-brain voxel-wise correlation maps obtained using the PF-fMRI superimposed to the standard MNI brain. The Pearson correlation (*r*) was computed voxel-wise across the whole brain, between the reconstructed activity inducing signals 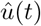 and the five motor tasks simulated as piece-wise constant signals with ones in the time points where the subject is executing the task and zeros elsewhere. The values *r* of the correlations are indicated by the color-bars. Each row corresponds to a specific motor task, from the top to the bottom: the tongue, the right and left hand, and the right and left foot. A: anterior; P: posterior; S:superior; I: inferior; R: right; L: left.

Quantitative comparison between the activity-inducing signal recovered using the PF-fMRI and the TA are shown in Figure 8. Results show that the mean Pearson correlations values estimated for each ROI across the data sample, increase while increasing the number of iterations, until it converges after 25 iterations for the hands, 5 iterations for the feet and about 35 iterations for the tongue. Moreover, starting from the first iteration, we show statistically significant higher correlation values compared to the ones obtained using the TA. In particular for the comparison between the PF-fMRI and the TA, Figure 9 shows the reconstructed signals 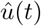 (Figure 9.a) and the correlations values (Figure 9.b) for a single subject (100307). We show a clear difference between the correlation values estimated in the area involved in the task and the one that is not involved. In fact, we show higher correlation between the tongue activation and the recovered activation 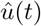 in the ROI rBA4p, that was expected to be involved in the motor task, while a low correlation is shown in the ROI rTE12 that instead is not involved. The TA was not able to clearly distinguish between an active and an inactive region since it showed low correlation values for both ROIs. As for the resting-state data analysis, in Figure 10 we show few exemplifying spatial maps over the 1200 analyzed using the PF-fMRI, between 215 and 350 TRs. The figure shows a dynamic between the regions composing the default mode network, i.e. the posterior cingulate corex, the medial prefrontal cortex, the lateral parietal lobules and the temporal cortex [53]. As shown in the figure, these areas are not all active simultaneously, but they assemble and disassemble over time in different combinations.

**Figure 8.**
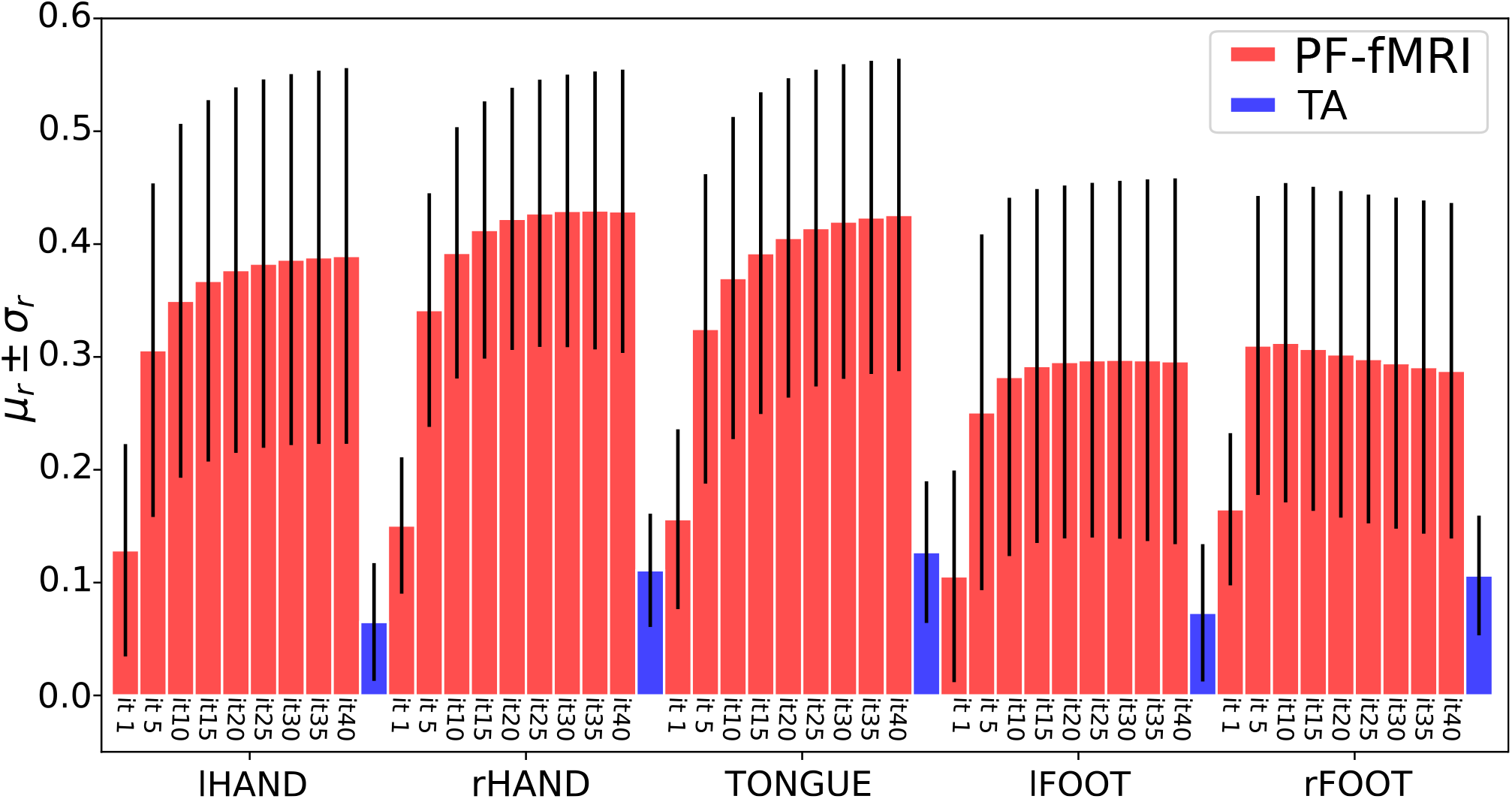
Barplots of the mean (*µ*_*r*_)±standard deviations (*σ*_*r*_) of the Pearson correlation coefficients (*r*) computed on the sample data (51 subjects) in 5 ROIs related to the tasks of the left and right hand, the tongue end the left and right foot. For each task, the bars in red represent the results using the PF-fMRI for an increasing number of iterations (from 1 to 40). The blue bar represents the activity-inducing signal estimated using the TA. (lHAND: left hand; rHAND: right hand; lFOOT: left foot; rFOOT: right foot.)

**Figure 9.**
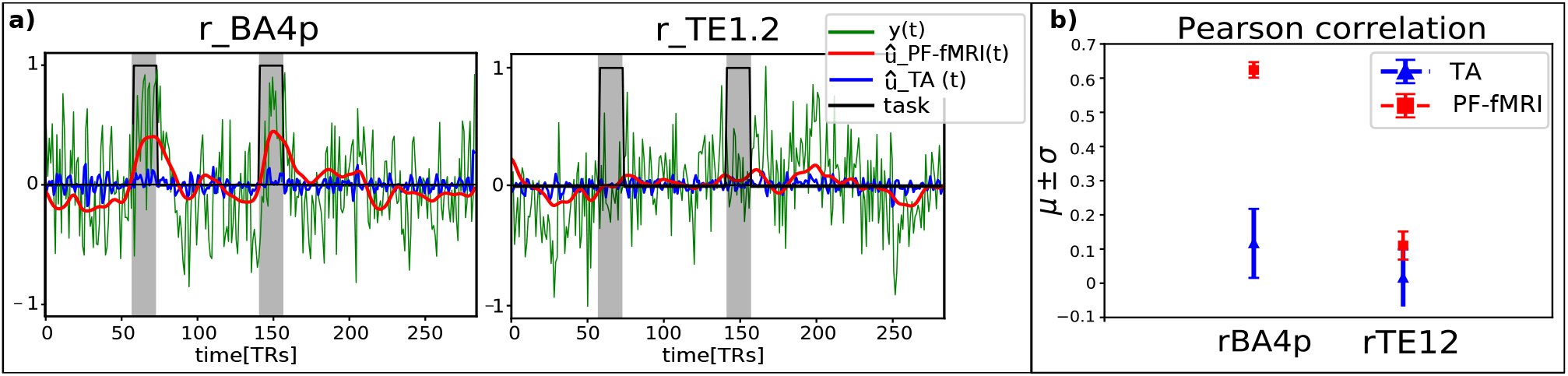
**a)** Reconstructed signals 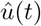 obtained with the PF-fMRI (red) and the TA tool (blue) superimposed on the real acquired fMRI signals (green) and the simulated tongue activation (black). The plot on the left is related to the ROI located on the Brodmann Area 4p (rBA4p), the plot on the right is associated to the ROI positioned on the primary auditory cortex (rTE1.2). All the signals were averaged across the voxels belonging to the GM-masked ROIs. The gray areas represent the occurrence and the duration of the tongue movements. **b)** Mean Pearson correlation coefficients (*µ*) and their associated standard deviations (*σ*) computed between the tongue activation and the recovered signals 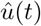 averaged across the voxels belonging to the GM-masked ROIs (rBA4p on the left, rTE1.2 on the right).

**Figure 10.**
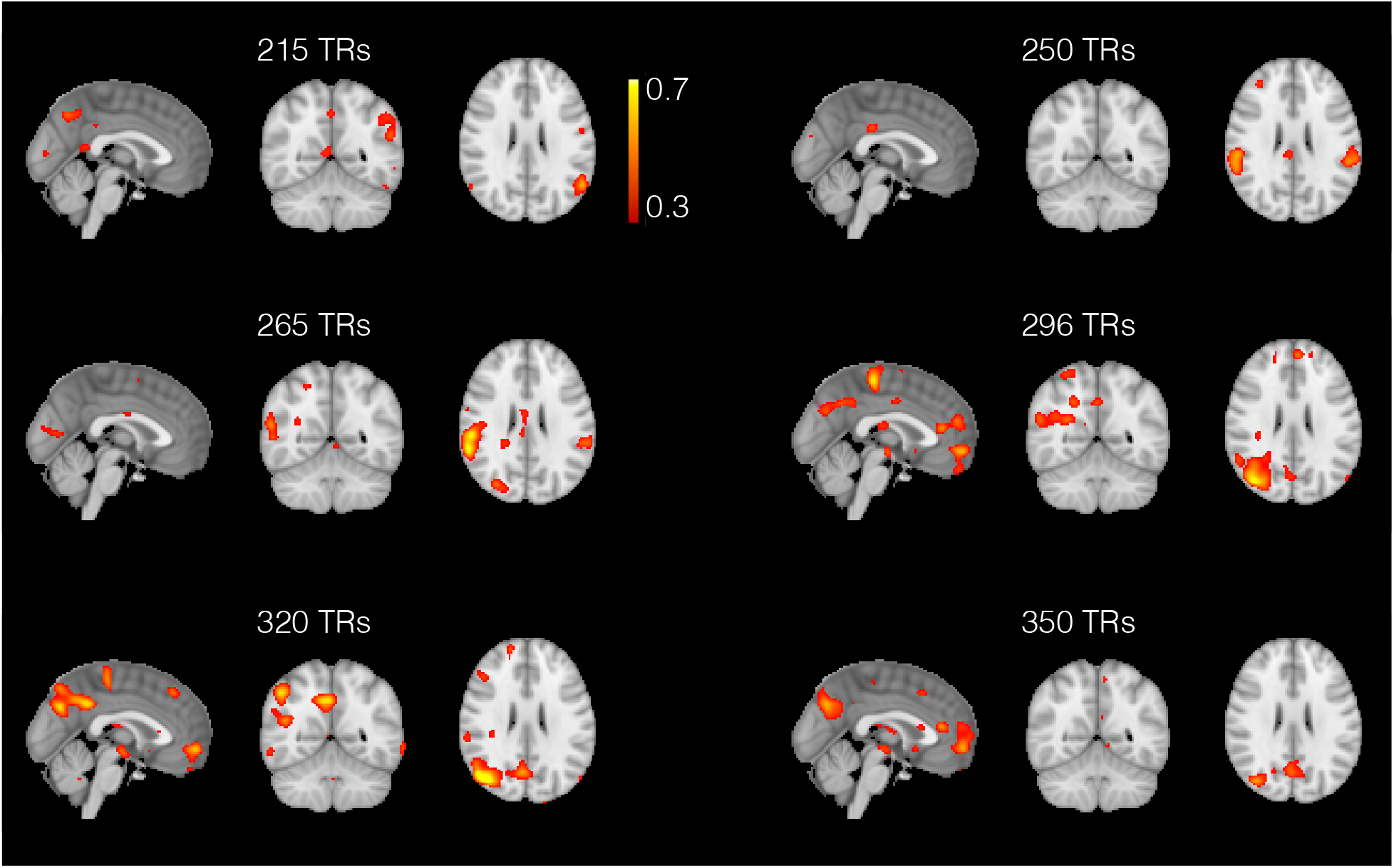
Examples of spatial maps obtained with the PF-fMRI spanned across 150 TRs (from 200 to 350 TRs). The maps shows the dynamics between the aread composing the default mode network. These regions are not all active at the same time but dynamically assemble and disassemble over time. The maps shows for example how the lateral parietal lobules are not always active simoultaneously as well as the medial prefrontal cortex that can activate selectively with the posterior cingulate cortex and the right temporal lobe.

## 5 Discussion

In this paper, we have described an innovative method to analyze fMRI images and recover the location and the occurrence in time of the functional neural activation. The proposed approach achieves this blindly, without the necessity of a priori knowledge of timing, duration and location of the underlying activation. The approach we proposed, namely the PF-fMRI, geometrically regularize the fMRI image such that: it saves and highlights large image variations as they are present at the occurrence of a brain activation or in the presence of a spatial edge with respect to small image variations, that instead are removed to reduce noise. To do this, we used the PDEs in an iterative algorithm and exploited the 4-D image structure tensor, that defines the directions of the gradient in the neighborhood of a voxel and directs towards an anisotropic or isotropic regularization. This gradient contains all the four principal directions of the fMRI image that is composed by a 3-D spatial image repeatedly acquired in time, which suggests that the whole 4-D fMRI image was smoothed contemporaneously in space and time at once.

Other approaches have been proposed to analyse fMRI data. Among them are the *(i)* the GLM, that fits a linear model to the fMRI time series, but it assumes prior knowledge of the tasks [1]; *(ii)* deconvolution methods, which are used to uncover brain activation from the BOLD response without prior information on the underlying activity [2, 15, 25, 27]. The deconvolution approach in [2] splits the problem into a spatial and temporal regularization problems, meaning that the user has to specify two regularization parameters and the two weights used to have a solution that is given by a weighted sum of the two separates regularization processes. In contrast, the PF-fMRI overcomes several limitations found in the previous literature. When comparing the regions recovered using the GLM and PF-fMRI, we noted overall very good agreement between the two methods. It should again be emphasized that, while the GLM requires knowledge of the experimental paradigm, the PF-fMRI does not. These results highlight that the PF-fMRI can be used to identify brain activity in the absence of an experimental paradigm, such as rs-fMRI. When comparing correlation maps obtained for the PF-fMRI and the TA, correlation values obtained with the PF-fMRI were significantly higher than those obtained with the TA suggesting an improved recovery of brain activity.

The PF-fMRI can be used for different purposes, for example to recover brain activation in a task experimental paradigm as well as in a rs-fMRI study, where the subject is asked not to perform any task while lying in the MRI scanner. Hence, the PF-fMRI could be useful to analyze the functional brain activity for those subject affected by neurological diseases who makes them unable to perform a task or to analyze unexpected brain activities, for example in the case of epilepsy, thus improving the recovery of brain dynamics for future clinical applications. The PF-fMRI could help in the recovery of time series and spatial maps that could be post-processed afterwards to perform statistical analyses. In particular, the recovered signals from rs-fMRI data could be further employed in the approach proposed by Karahanoğlu and Van De Ville [54] to investigate the dynamics of resting-state networks and to reveal transients in spontaneous neural activity. Results, and specifically the ones shown in Figure 10, show the great potentials of the PF-fMRI to analyze the dynamics of rs-fMRI data [55] on a larger sample in healthy control as well as in a patient group. The PF-fMRI could also be exploited in a framework that employs the dynamic causal modeling [56] or those that try to infer white matter information flow [57]. It must be clarified that the PF-fMRI does not any hypothesis on the interactions between brain regions. Once the activation is inferred using the PF-fMRI, the dynamic causal modelling could be expolited on the recovered signals to reveal possible causality between brain regions’ activity. Also, it had been shown that including fMRI information from the PF-fMRI approach to find active regions to be exploited as priors on cortical regions allows to select plausible structural connections to yield a tractable optimization problem [57]. The information flow between cortical regions known to be active can be blindly identified with the PF-fMRI and without requiring a manual selection of the regions of interest. One limitation of the PF-fMRI is that it does not generate a very sharp piece-wise constant function, but instead produces a smooth version of the activity-inducing signal. Future works could include into the solution of the inverse problem the information given by the diffusion MRI data that would provide us with a more complex neighborhood defined by white matter connectivity. In this way, the neighborhood would no longer be given only by the surrounding voxels, but also by the voxels which are anatomically segregated and therefore functionally connected to achieve the same function. In addition, the proposed PF-fMRI approach could be exploited to investigate possible fMRI activation in the white matter, that is an emerging debated topic in the neuroimaging field [58].

## 6 Conclusion

In this paper we proposed and validated a new method to blindly regularize the fMRI images and recover the brain activity from fMRI signals without prior knowledge. Our findings show that the PF-fMRI enabled us to solve an important problem, coupling the spatial and the temporal dimension, and to recover brain activation overlapping the ones obtained with the GLM. Our results also show higher correlations of the recovered time courses with the ground truth compared to the TA. This opens a new channel for the analyses of rs-fMRI data and the recovery of paradigm free neural activity, to be used for investigations in future clinical applications.

## Acknowledgments

This work has received funding from the European Research Council (ERC) under the European Union’s Horizon 2020 research and innovation program (ERC Advanced Grant agreement No 694665: CoBCoM -Computational Brain Connectivity Mapping).

Data were provided [in part] by the Human Connectome Project, WU-Minn Consortium (Principal Investigators: David Van Essen and Kamil Ugurbil; 1U54MH091657) funded by the 16 NIH Institutes and Centers that support the NIH Blueprint for Neuroscience Research; and by the McDonnell Center for Systems Neuroscience at Washington University.

## Footnotes

1 https://fsl.fmrib.ox.ac.uk/fsl/fslwiki

2 https://fsl.fmrib.ox.ac.uk/fsl/fslwiki/POSSUM

3 https://miplab.epfl.ch/index.php/software/total-activation

